# Scalable Fabrication of a Tough and Recyclable Spore-Bearing Biocomposite Thermoplastic Polyurethane

**DOI:** 10.1101/2024.10.25.620092

**Authors:** Han Sol Kim, Evan M. White, Grant Crane, Kush Patel, Myung Hyun Noh, Md Arifur Rahman, Adam M. Feist, Jason J. Locklin, Jonathan K. Pokorski

**Affiliations:** Aiiso Yufeng Li Family Department of Chemical and NanoEngineering, University of California San Diego, 9500 Gilman Dr., La Jolla, CA 92093, USA; New Materials Institute, University of Georgia, Athens, GA 30602, USA; Shu Chien-Gene Lay Department of Bioengineering, University of California San Diego, 9500 Gilman Dr., La Jolla, CA 92093, USA; Thermoplastic Polyurethane Research, BASF Corporation, 1609 Biddle Ave., Wyandotte, MI 48192, USA; Institute for Materials Discovery and Design, University of California San Diego, 9500 Gilman Dr., La Jolla, CA, 92093, USA

**Keywords:** Polyurethane, Biocomposite polymer, Spore, Toughness, Scale up, Recycling

## Abstract

Thermoplastic polyurethanes (TPUs) are a class of versatile thermoplastic elastomers where most of their products lack a proper recycling strategy or have no end-of-life solutions. To pursue a sustainable end-of-life solution for TPU based products, self-disintegrating biocomposite TPUs have recently been developed by embedding spores of TPU-degrading bacteria into TPUs using melt extrusion. Herein, we improve upon spore-bearing biocomposites and demonstrate industrially-relevant manufacturing conditions for the fabrication of biocomposite TPUs. Spore production was modified to reduce the innate brown color of the resulting materials and, effectively minimize the coloration of biocomposite TPUs. Reduction of FeSO_4_ in a sporulation media generated white spores without compromising spore productivity, viability, morphology or heat-shock tolerance. Biocomposite TPUs containing white spores displayed a 45% increase in toughness compared to TPUs without spores, while retaining ∼90% spore viability post processing. Furthermore, biocomposite TPU fabrication was demonstrated using a scalable continuous extruder followed by injection molding. Biocomposite TPUs generated by these industry-relevant processes exhibited comparable toughness improvement and spore viability to biocomposite TPU prepared using a lab scale microcompounder, while enhancing productivity by 30-fold. Finally, spore addition improved biocomposite TPUs recyclability, enabling 80% toughness retention after 5 rounds of iterative melt processing. Additionally, no negative effect on the lifespan of the generated TPUs was observed over a duration of 1 year of storage. Overall, this study confirmed that spore-bearing biocomposite TPUs are promising for practical applications, offering an accessible method to enhance toughness and sustainability of commercial TPUs through the incorporation of spore-based living fillers.

## 1. Introduction

Thermoplastic polyurethanes (TPUs) are among the most versatile elastomers extensively used in products such as phone cases, footwear and automotive parts. TPUs are synthesized by the reaction of polyols, isocyanates, and chain extenders, generating block copolymers with good thermal and mechanical properties[1]. Polyol blocks in TPUs provide a soft matrix, while the adducts of isocyanates and chain extenders form hard domains based on self-assembly promoted by hydrogen bonding, π-π stacking, and/or van der Waals’ interactions[2–4]. The combination of alternating soft and hard segments, together with the availability of diverse TPU building blocks, modulates unique intra- and interchain interactions in TPUs, rendering their outstanding and tailorable properties[5]. Consequently, polyurethanes display desirable properties, such as flexibility, rigidity, and abrasion resistance, and have been the sixth-most produced polymer globally with growing demands[6].

However, TPUs possess environmental issues at their end-of-life due to the lack of commercially available recycling streams. Polyurethanes can be collected under category 7 of the resin identification code for miscellaneous plastics, but only 0.3% of plastics collected in this category are recycled in the US[7]. Instead, most TPU-based products are landfilled, leached into the environment as uncontrolled waste, or incinerated after their useful life[8]. To mitigate accumulation of TPU waste, we recently reported self-disintegrating biocomposite TPUs, in which spores of TPU-degrading bacteria (i.e., an evolutionarily engineered *Bacillus subtilis* ATCC 6633 strain, ATCC 6633 HST) were embedded within the TPU matrix as living fillers[9]. Spores embedded within TPUs served as polymer reinforcing agents and resulted in up to a 37% improvement in toughness. At the end of the material’s life, spores could be germinated to facilitate the disintegration of the TPU. The biocomposite TPUs showed great promise as a robust and sustainable TPU alternative on a laboratory scale, however practical implementation of spore-bearing TPUs will have several challenges beyond the laboratory scale, including filler quality and cost, as well as scale-up to commercial equipment.

Herein, we explored the feasibility of spore-bearing biocomposite TPUs in various aspects and demonstrated their scalable fabrication (**Scheme 1**). Reducing FeSO_4_ concentration in the sporulation media produced white spores without affecting their viability, productivity, or morphology, effectively suppressing the undesirable brown coloration in spore-bearing TPUs, while maintaining a visual appearance similar to virgin TPU. Spores compounded with TPU not only served as TPU reinforcing fillers, improving toughness of the TPU by up to 45%, but also as radical scavenging agents enabling reprocessing with minimal deterioration of mechanical properties. Finally, the scalable fabrication of biocomposite TPUs using industry-relevant continuous extrusion and injection molding processes are demonstrated, as well as the long-term storage stability of biocomposite TPUs.

**Scheme 1.**
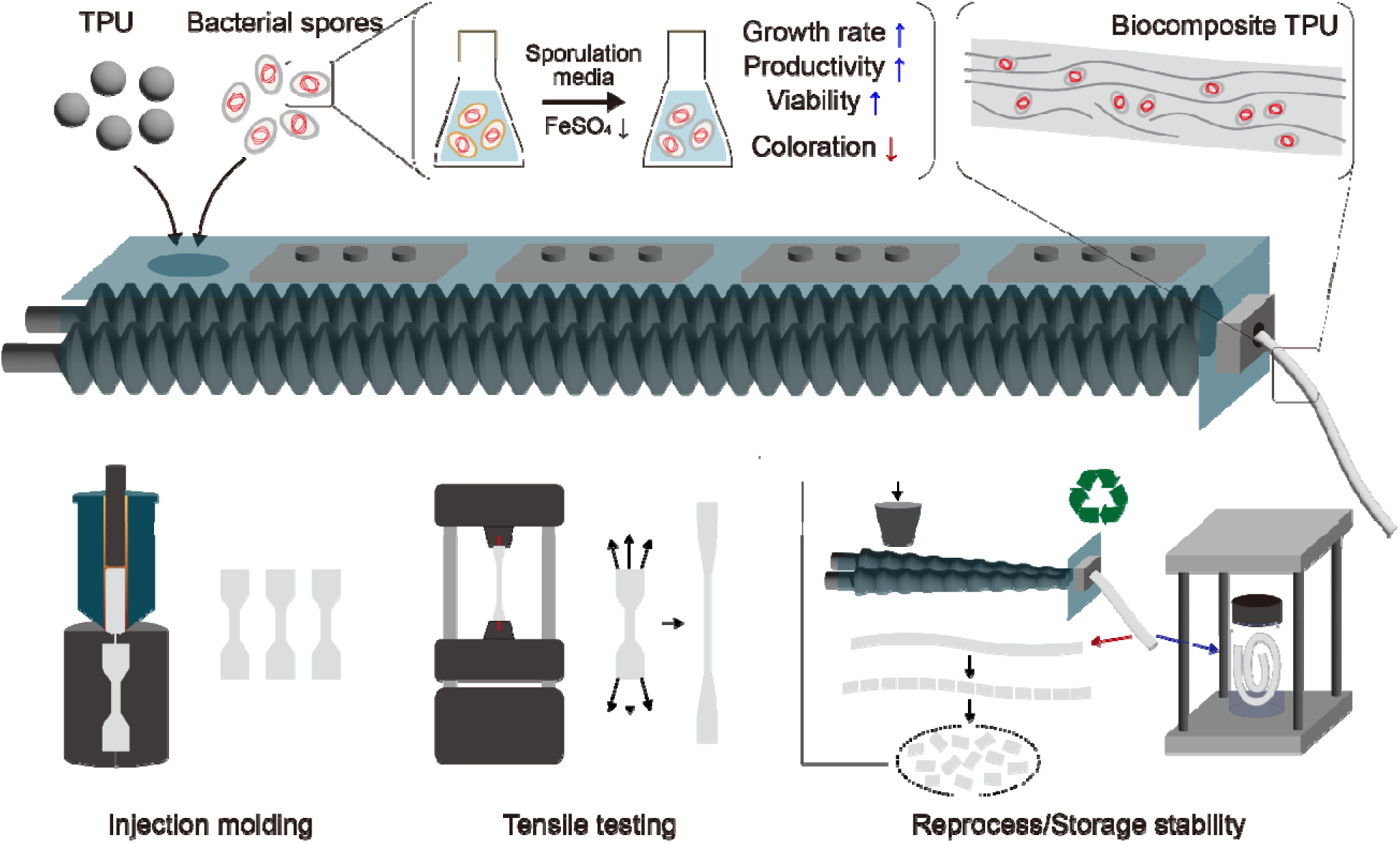
Scalable fabrication of spore-bearing biocomposite thermoplastic polyurethane (TPU).

## 2. Results and Discussion

### 2.1. Media modification to tune the color of generated spores

Color is an important component for plastic appearance and marketability. Bacterial spores prepared by previously reported protocols, however, inherently have a brown color and behave as colorants to composites after compounding with TPUs[9]. To mitigate the undesirable coloration of biocomposite TPUs, we developed a new spore production protocol. The brown color of the spores could be attributed to pulcherriminic acid secreted by *B. subtilis* during sporulation, which binds with iron to form iron chelate pulcherrimin as a brown pigment[10,11]. Consistent with this previous work, we found that FeSO_4_, a metal salt added to the originally utilized sporulation media, was responsible for the brown color in the final lyophilized spore powder. We reduced the concentration of FeSO_4_ by 1000 times (from 1 µM to 1 nM) from our previously-reported sporulation protocol [9] and found that the color of the purified and lyophilized spores was significantly reduced from brown to white (**Fig. 1**). The reduction of FeSO_4_ concentration in the sporulation media not only reduced the color of spores (**Fig 1A**), but also enhanced spore biomass productivity (mass of spores produced per liter culture per day, after lyophilization, **Fig. 1B**) and specific spore viability (colony forming units per mg spore powder, **Fig. 1C**) by 184% and 404%, respectively. Moreover, there was no morphological difference between spores prepared by using sporulation media with 1 µM or 1 nM FeSO_4_ (**Fig. 1D**). The survivability of the “white” spores after the heat-shock treatment at 100 °C for 10 min was also not compromised, showing a comparable heat-shock tolerance with the original “brown” spores (**Fig. 1E**). Even though iron is known to be required for the sporulation of *B. subtilis*[12–14], several studies have shown that excessive amount of divalent cations in sporulation media, including Fe^2+^, can decrease spore production yield[15]. ATCC 6633 HST strain positively responded to the reduction of FeSO_4_ concentration, resulting in an overall enhanced spore productivity and viability. Furthermore, spores produced using modified sporulation media retained their heat-shock tolerance and, importantly, presented a white color that can potentially reduce the coloration of spore-bearing TPU composites.

**Fig. 1.**
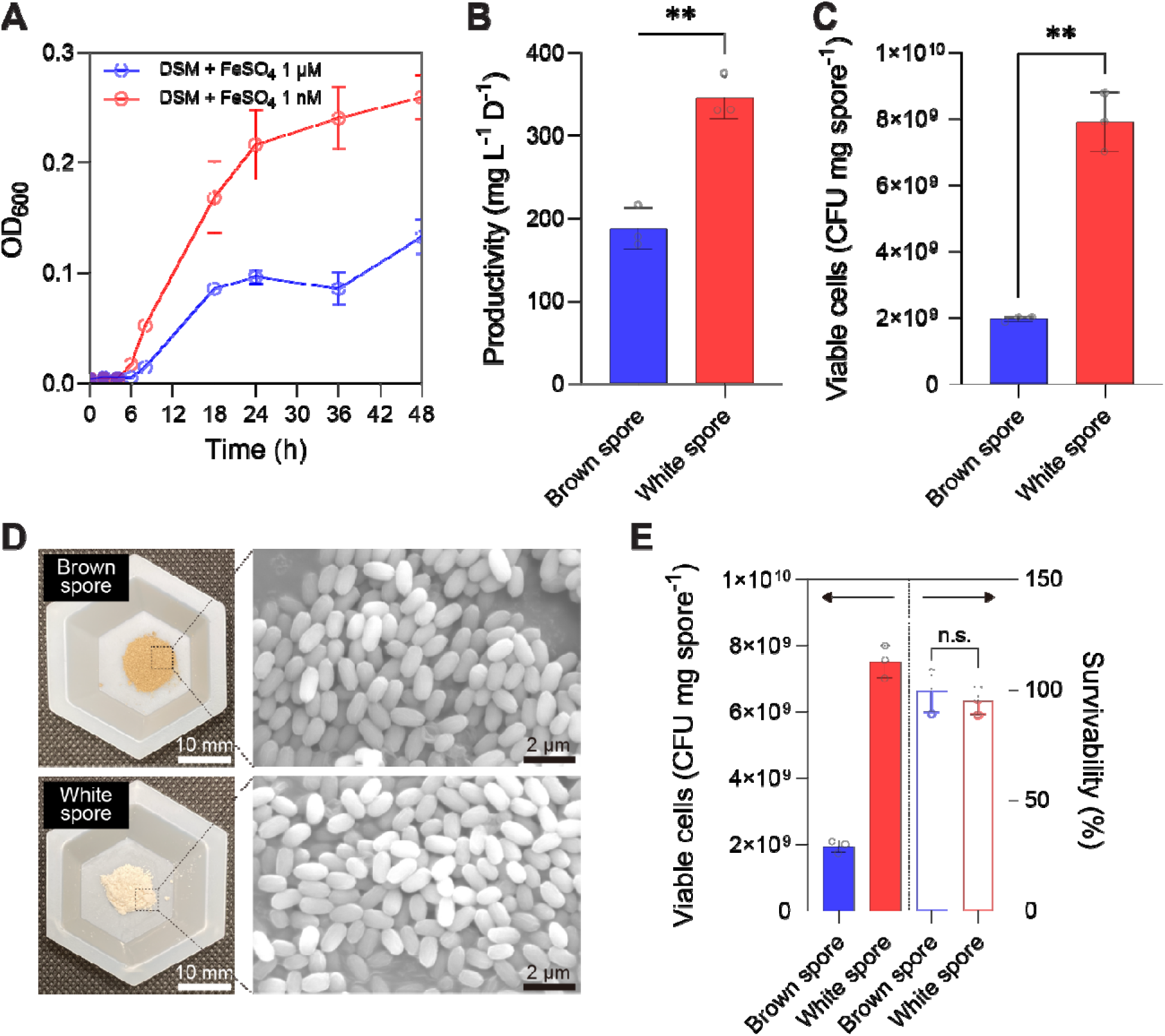
Impact of sporulation media modification. (A) OD_600_ profiles of B. subtilis cultured/sporulated in sporulation media with two different FeSO_4_ concentrations (1 µM and 1 nM). (B) Productivity in dry weight and (C) specific viability of lyophilized spores. (D) Photographs (scale bars: 10 mm) and scanning electron microscope images (scale bars: 2 µm) of spores. (E) Specific viability and survivability of spores after heat-shock treatment unde 100 °C dry heat for 10 min. Statistical analysis was performed using unpaired t-test (n = 3 pe group; n.s. = not significant (P > 0.05); **P < 0.01)

### 2.2. Tensile properties of biocomposite TPU fabricated by utilizing white spores

Biocomposite TPUs with brown (BC TPU^Brown^) or white (BC TPU^White^) spores were prepared at different spore loading levels from 0.00% (w/w) to 1.00% (w/w) (**Fig. 2A & 2D and Fig. S1**) and their tensile properties were evaluated. Utilization of white spores significantly reduced the coloration of BC TPUs, showing minimal color change when compared to TPU without spores. BC TPU^Brown^ exhibited a very consistent toughness profile with our previous report, showing up to a 39% toughness improvement upon the addition of 0.75% (w/w) spores. However, BC TPU^White^ showed a greater toughness improvement (up to 45%) at a lower spore loading (0.50% (w/w)) compared to BC TPU^Brown^ (**Fig. 2B & 2E**). Significantly improved toughness by spore incorporation suggested that spores behaved as polymer reinforcing fillers based on interfacial adhesion between the spore surface and TPU matrix[9,16]. Decreased toughness enhancement above critical spore loadings (0.75% (w/w) and 0.50% (w/w) for BC TPU^Brown^ and BC TPU^White^, respectively) can be explained by the aggregation of spores in TPU matrix. Spores at higher concentration in a low viscosity TPU melt tend to aggregate due to mutual attraction among spores and lead to non-homogeneous mixing of TPU and spores[17]. Such non-homogeneous mixing of spores disrupts proper load bearing at the interface of TPU and spores which leads to earlier failure in tension.

**Fig. 2.**
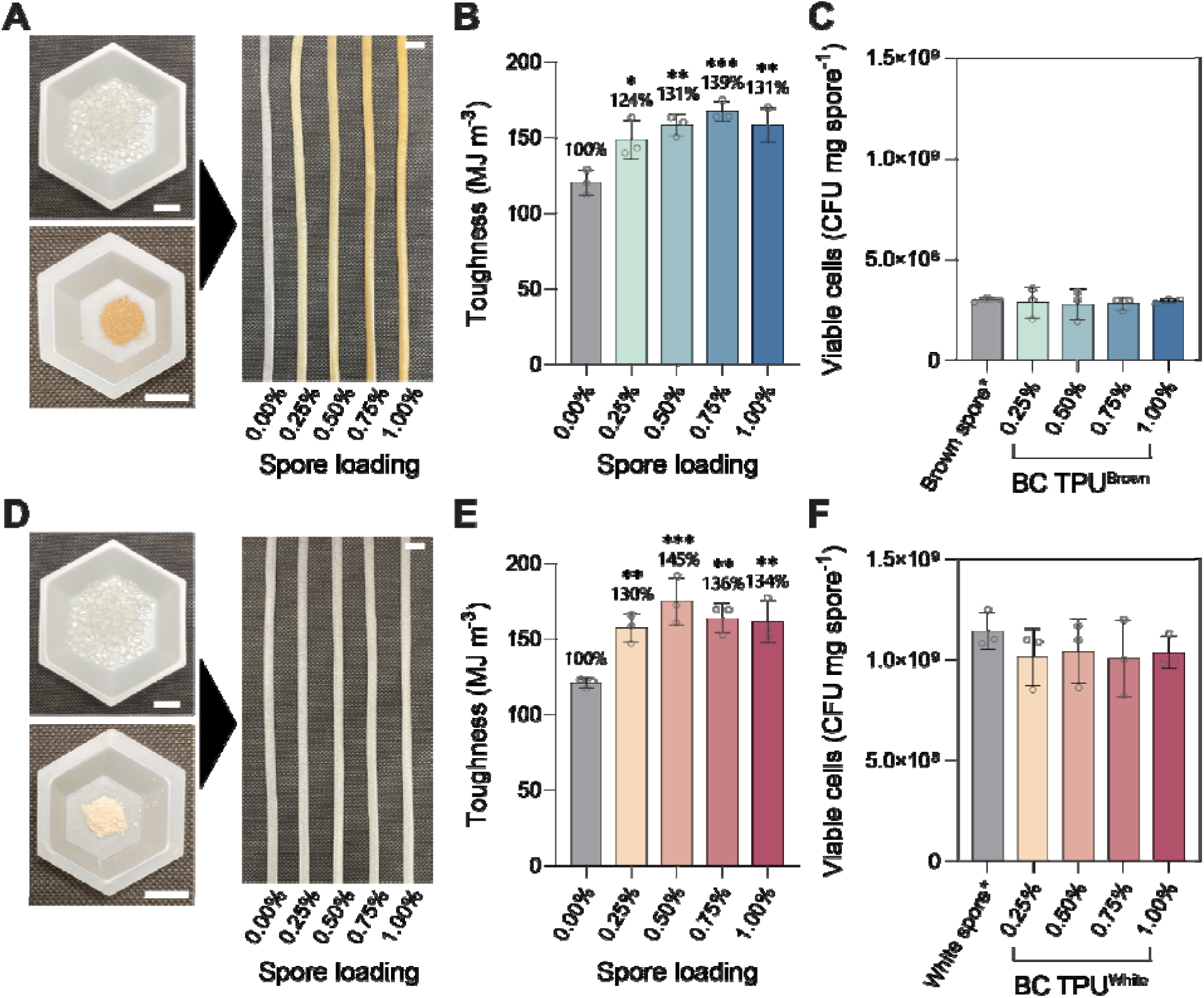
Fabrication of spore-bearing biocomposite TPUs (BC TPUs). Photographs of TPU pellets (top left panels), spore powder (bottom left panels) and BC TPU strips (right panels) (A&D), and characterization of their tensile toughness (B&E) and spore survivability (C&F). Biocomposite TPUs were fabricated by using either brown (A-C) or white spores (D-F) at 0.00 ∼ 1.00% loading. Scale bars: 10 mm. One-way analysis of variance (ANOVA), followed by a post-hoc test with two-sided Dunnett’s multiple comparisons, was used for statistical comparison between TPU (control) and BC TPUs (n = 3 per group; *P < 0.05; **P < 0.01; ***P < 0.001;)

To assess biological activity incorporated into TPU, spores inside BC TPUs were extracted by solvent treatment, followed by quantification of viable spores via CFU assay. Specific viability of spores extracted from BC TPUs were compared with that of spores that followed the same solvent treatment procedures to evaluate spore survivability. After compounding with TPU, brown and white spores in BC TPUs retained ∼95% and ∼90% survivability, respectively, regardless of spore loading (**Fig. 2C & 2F**). It should be noted that BC TPU^White^ displayed ∼3.6-fold higher specific spore viability compared to BC TPU^Brown^ (**Fig. 2C & 2F**), counterbalancing the slightly reduced spore survivability in terms of number of active spores in composites. Given that these advances were achieved by simply changing FeSO_4_ concentration in the sporulation media, white spores can be feasible biofillers for biocomposite TPU fabrication.

Other tensile properties of BC TPU^Brown^ and BC TPU^White^, such as ultimate tensile stress, elongation at break and Young’s modulus are presented in **Fig. S2**. Both brown and white spores improved the ultimate tensile stress and elongation at break, both of which contributed to the improved toughness of BC TPUs. Spore addition did not significantly affect the Young’s modulus of the TPUs. White spores were used for further experiments unless otherwise noted.

### 2.3. Continuous manufacturing and injection molding of biocomposite TPU

We evaluated the feasibility of biocomposite TPU fabrication using continuous extrusion followed by injection molding (**Fig. S3**). The continuous extruder we utilized in this study (Thermo Fisher Process 11) adapts parallel twin screws with a 40:1 L/D ratio, replicating a viable method for scale-up by simulating intensive compounding and high-speed processing[18]. Continuous processing enabled a remarkable improvement in both the rate and yield of biocomposite TPU fabrication. For example, a control batch mode processing run mimicking our original demonstration[9] in a laboratory-scale extruder generated 7.5 g h^-1^ BC TPUs with 50 ∼ 60% sample loss, primarily as residues in the cycling channel (final productivity: 3.3 g h^-1^). However, continuous manufacturing showed >100 g h^-1^ throughput with minimal sample loss during processing, leading to 30-fold productivity improvement (**Fig. 3A**). Importantly, extrusion conditions optimized in the batch mode extruder (135 °C barrel temperature / 36 rpm screw speed) were successfully transferred to continuous operation with minor modifications (135 °C barrel temperature / 50 rpm screw speed). BC TPUs manufactured by continuous extrusion were subsequently processed into standard dog bone shapes using injection molding for tensile testing (ASTM D638 Type V).

**Fig. 3.**
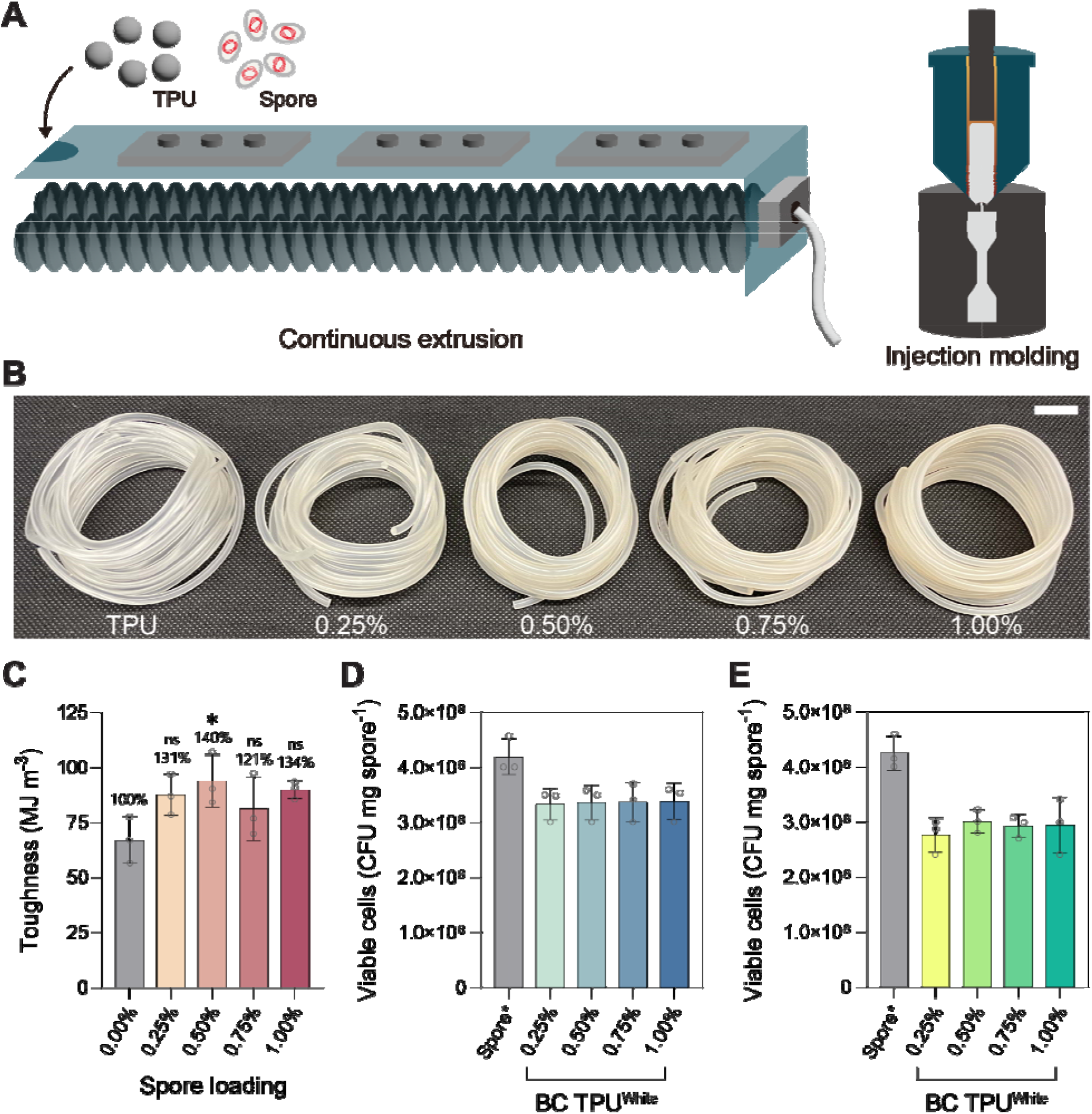
Scale up of biocomposite TPU fabrication. (A) Schematic illustration of sequential continuous extrusion and injection molding. (B) Biocomposite TPU with 0.00 ∼ 1.00% spore loading fabricated by continuous extruder. Scale bar: 20 mm. (C) Toughness of biocomposite TPU sequentially processed by continuous extrusion and injection molding. Spore survivability post continuous extrusion (D) and subsequent injection molding (E). One-way ANOVA, followed by a post-hoc test with two-sided Dunnett’s multiple comparisons, was used for statistical comparison between TPU (control) and BC TPUs (n = 3 per group; ns = not significant, *P < 0.05)

BC TPU^White^ exhibited up to 40% toughness improvement at 0.5% (w/w) spore loading even after being processed by the combination of continuous extrusion and injection molding (**Fig. 3B**). Such strong consistency of toughness improvement between BC TPUs prepared and characterized by different practices confirmed that spores were reliable reinforcing agents for TPU. Spore survivability after continuous extrusion and injection molding was 80% and 70%, respectively (**Fig. 3C-D**). The continuous extruder generated 23 GPa shear stress (50 rpm screw speed), which is approximately 250-fold higher than that of the maximum shear stress generated by the batch mode extruder (90 MPa)[9]. At the same time, biocomposite TPUs were subjected to high heat (140 °C) and pressure (300 ∼ 600 bar) during injection molding. 70% spore survivability under such rigorous conditions implied that spores were inherently resistant against the applied shear stress. Other tensile properties of BC TPU^White^ sequentially processed by continuous extrusion and injection molding are presented in **Fig. S4**. Similar to lab scale demonstration, spore addition improved both the ultimate tensile stress and elongation at break, while BC TPUs remained as soft as TPU without spores by showing a similar Young’s modulus.

### 2.4. Reprocessing by extrusion and long-term storage of biocomposite TPU

Polymer processing at high temperature generally induces thermal degradation of polymers by the supply of external heat and/or by the internally-generated heat through dissipation[19]. Such thermo-mechanical and/or thermo-oxidative degradation of polymers have been great challenges for polymer recycling[19]. We hypothesized that the bacterial spores embedded within the TPU would serve as an antioxidant and prevent degradation upon reprocessing/thermo-mechanical recycling. To evaluate this, the tensile property change of reprocessed TPU by hot melt extrusion (HME) was measured after 5 rounds of consecutive compounding to simulate TPU recycling. Virgin TPU showed a gradual toughness decay with increasing iterations of reprocessing (**Fig. 4A**). Freshly processed TPU via HME (cycle 1) exhibited 134.4 MJ m^-3^ toughness, whereas 4-times reprocessed TPU (cycle 5) presented 67% residual toughness (88.8 MJ m^-3^). On the other hand, BC TPU^White^ with 0.5% (w/w) spores before (cycle 1) and after 4 rounds of reprocessing (cycle 5) showed 185.2 MJ m^-3^ and 161.7 MJ m^-3^ toughness respectively, retaining 87% of toughness after multiple extrusion (**Fig. 4B**). Interestingly, after the initial toughness loss (cycle 2), BC TPU^White^ maintained 87 ∼ 89% toughness during the subsequent reprocessing (cycles 3 - 5) (**Fig. 4B**). Recyclability of TPU materials could be further illustrated by the color change observed in reprocessed extrudates. TPU without spores exhibited increasing yellowing after multiple extrusion cycles, whereas BC TPU^White^ did not exhibit a significant color change (**Fig. 4A-B**)[20].

**Fig. 4.**
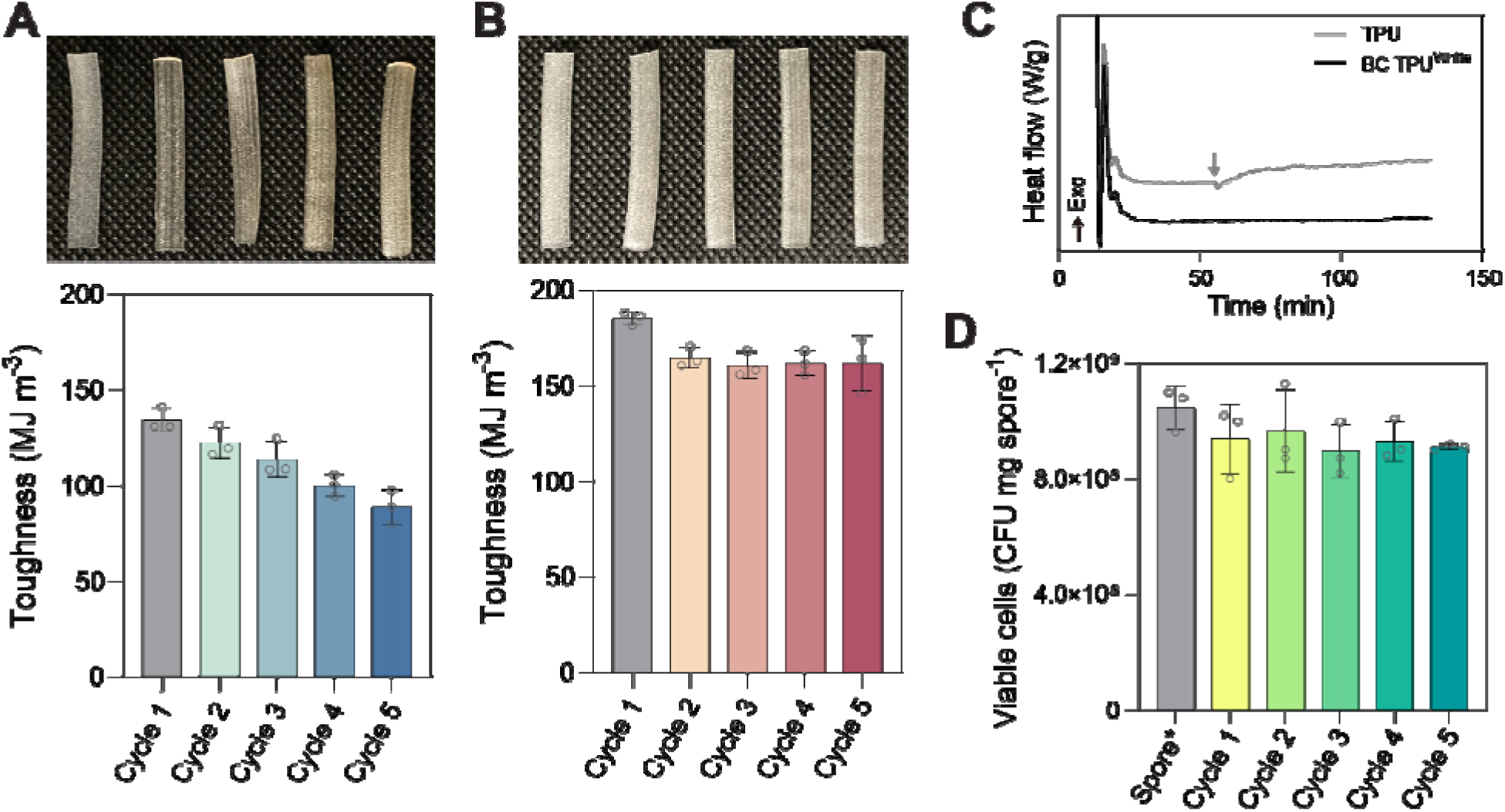
Reprocessing TPU and biocomposite TPU to simulate polymer recycling. Photographs (top) and toughness (bottom) of reprocessed TPU (A) and BC TPU (B) for multiple times (scale bars: 10 mm). (C) DSC under O_2_ environment to determine OIT. Grey arrow indicates the onset of thermal degradation of TPU. (D) Spore survivability in biocomposite TPU during recycled processing.

To investigate the mechanism behind the recyclability of spore-bearing TPU, differential scanning calorimetry (DSC) was performed under an oxygen environment to analyze oxidative induction time (OIT). After 43 min of isothermal treatment at 150 °C under 100% O_2_ environment, thermal degradation of TPU without spores was initiated. In contrast, no thermal degradation was found from BC TPU^White^ under the same condition for 2 h (**Fig. 4C**). This finding suggested that spores in the biocomposite TPU not only served as polymer reinforcement that improved the toughness by ∼40%, but also functioned as an antioxidant that prevented thermal degradation of the polymer during melt processing by scavenging radicals generated by internal/external heat[21]. Strong interfacial adhesion between the spore additives and TPU could have been another contributor to the recovery of tensile properties after the thermomechanical recycling of BC TPUs[22]. Other tensile properties of reprocessed TPU and BC TPU^White^ are presented in **Fig. S5**. Both ultimate tensile stress and elongation at break of TPU decreased with each reprocessing cycle in the absence of spores. In contrast, BC TPU^White^ retained its full elongation at break throughout 4 reprocessing cycles. Ultimate tensile stress of BC TPU^White^ was maintained at ∼90% level after the initial drop. Reprocessing had no significant impact on the Young’s modulus of either TPU or BC TPU^White^. Spores in BC TPU^White^ retained ∼90% spore survivability post 4 rounds of reprocessing (**Fig. 4D**).

Further, we evaluated the long-term storage stability of TPU and biocomposite TPU. TPU without spores and BC TPU^White^ (0.5% (w/w)) were stored under ambient conditions (23 °C and 60 % relative humidity) for up to 1 year. Both TPU and BC TPU^White^ maintained their tensil properties over the 1 year of shelf storage, suggesting that spore addition has no negative effect on the lifetime of TPU (**Fig. 5A-B & Fig. S6**). In addition, spores in BC TPU^White^ remained viable for 1 year (**Fig. 5C**).

**Fig. 5.**
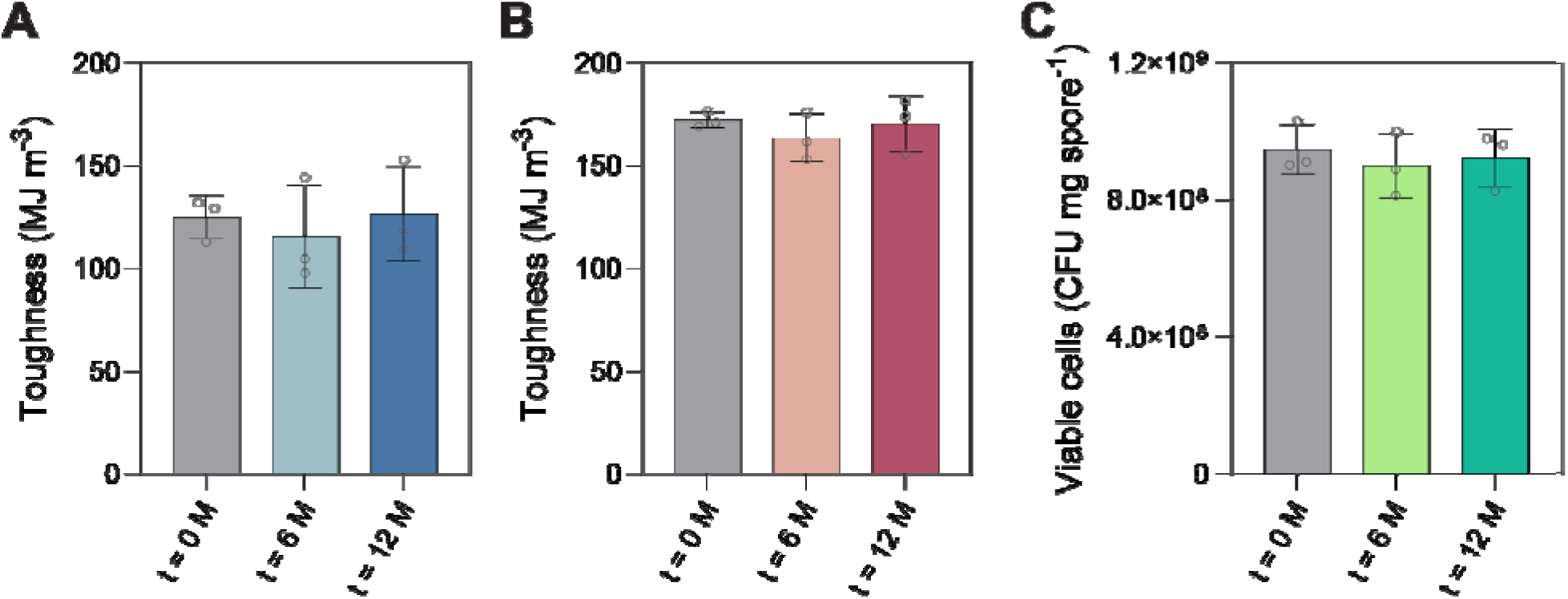
Biocomposite TPU longevity analysis. Toughness of TPU (A) and biocomposite TPU (B) during 1 year storage under ambient conditions. (C) Spore survivability in biocomposite TPU during shelf storage.

## 3. Conclusions

In this study, an accessible and scalable method to produce spore-bearing biocomposite TPUs was demonstrated. An undesirable coloration of TPU by spores was resolved by modifying the FeSO_4_ concentration of sporulation media, which resulted in the generation of white spore without compromising spore productivity, viability, morphology or heat-shock tolerance. Biocomposite TPUs fabricated by utilizing the white spores showed 45% toughness improvement compared to virgin TPU without spores. Furthermore, the BC TPU^White^ exhibited maximum toughness improvements at lower spore loading (0.50% (w/w)) compared to the critical spore concentration of brown spores in TPU (0.75% (w/w)). By collectively considering enhanced toughness improvement (i.e., 39% increased to 45%) at a diminished spore loading (i.e., 0.75 % decreased to 0.50 %), we can hypothesize that the white spores can be a more effective TPU reinforcing filler compared to the brown spores. Notably, we demonstrated the scale-up of a biocomposite TPU fabrication by using a continuous extruder, which adopts a screw design geometrically similar to pilot or industry scale extruders and, thus, offers reliable scalability[23]. We also further challenged biocomposite TPU extrudate through injection molding to simulate the current practice of polymer processing. Biocomposite TPUs sequentially processed by continuous extrusion and injection molding displayed significant toughness improvement (up to 40%) and spore survivability (70%), confirming the compatibility of biocomposite TPU fabrication with industry-relevant processing. Further, spores in biocomposite TPU likely behaved as antioxidants scavenging free radicals generated during TPU compounding and, thereby, enabled the recycled melt processing of TPU material. Finally, spore addition had no negative effect on the lifetime of TPU during long-term storage for 1 year. Overall, biocomposite TPU showed excellent feasibility in terms of tensile properties, scalability, recyclability and longevity.

## 4. Methods

### 4.1. Bacterial cells and materials

Evolutionarily-engineered *Bacillus subtilis* strain developed in our previous work (A5_F40_I1 strain, ATCC 6633 HST) was used as spore-forming bacteria[9]. Soft-grade, polyester-based thermoplastic polyurethane pellets (Elastollan® BCF45) were gifted from BASF. Luria–Bertani (LB) powder (Miller), BD Difco^TM^ nutrient broth, BD Bactor^TM^ agar and 10x phosphate-buffered saline (PBS) pH 7.4 were purchased from Fisher Scientific (Waltham, MA, USA). KCl, MgSO_4_, CaCl_2_, FeSO_4_, MnCl_2_, lysozyme and N,N-dimethylformamide (DMF) were purchased from Sigma–Aldrich (St. Louis, MO, USA).

### 4.2. Bacteria culture and spore production

We followed our previously-reported protocols for the cultivation, sporulation and purification of bacterial spores with minor modifications[9]. In detail, LB medium and Difco sporulation medium (DSM) were used for seed and main culture of ATCC 6633 spore production, respectively. LB medium was prepared by dissolving 25 g L^-1^ LB powder in deionized (DI) water, followed by autoclaving at 121 °C for 20 min. DSM was prepared by dissolving 8 g L^-1^ BD Difco^TM^ nutrient broth, 1 g L^-1^ KCl and 0.12 g L^-1^ MgSO_4_ in deionized (DI) water. After autoclaving at 121 °C for 20 min, CaCl_2_ and MnCl_2_ were added to a final concentration of 1 mM and 1 µM, respectively. Additionally, 1 µM or 1 nM FeSO_4_ was added to the autoclaved DSM to produce brown or white spores, respectively.

Seed culture was prepared by inoculating ATCC 6633 HST into LB medium at 1 % (v/v) final concentration. The seed culture was incubated at 37 °C at 250 rpm shaking overnight. The seed culture was then inoculated into DSM at 1 % (v/v) final concentration for the main culture. The main culture was incubated at 37 °C at 250 rpm shaking for 2 days to ensure enough sporulation of ATCC 6633 HST.

Spores were collected from the liquid by centrifugation. Spore suspension was centrifuged at 12,000 g at 4 °C for 20 min (Avanti J-E, Beckman Coulter Life Science, Brea, CA, USA). After removing supernatant, recovered spore pellets were resuspended in 100 mM PBS pH 7.4. Spores were further washed with PBS by repeating the following steps for three times: (i) centrifugation at 2200 g at 25 °C for 10 min (5810 R, Eppendorf), (ii) supernatant removal, and (iii) suspension in 100 mM PBS pH 7.4. After three times of washing, spores were treated with 2.5 mg mL^-1^ lysozyme solution prepared in 100 mM PBS pH 7.4 to degrade remaining vegetative cells. After 1 h incubation at 37 °C at 250 rpm, spores were washed with PBS for three times. Spore suspension was then heat treated at 65 °C for 1 h under stationary conditions. Finally, the spores were washed with DI water three times. After freezing purified spore suspension using liquid nitrogen, spore suspension was lyophilized for 3 days using FreeZone 2.5 (Labconco, Kansas City, MO, USA).

### 4.3. Field-emission scanning electron microscopy (SEM)

Spore powder was loaded onto carbon tape, followed by gold coating for 1 min three times. Spores were visualized using FEI Apreo 2 SEM (Thermo Fisher Scientific, Waltham, MA, USA) at 10 kV at 3.2 nA.

### 4.4. Hot melt extrusion using batch extruder

5 g of TPU pellets was added to the lab-scale twin screw extruder, HAAKE^TM^ Mini CTW (Thermo Fisher Scientific) equipped with a slit exit die (slit size: 5.0 mm x 0.7 mm). TPU pellets were melted at 135 °C at 36 rpm screw speed for 5 min. Subsequently, spore powder was added to the extruder at 0 ∼ 1% (w/w) loading. TPU-spore mixture was compounded at 135 °C at 36 rpm screw speed for 15 min. The composite was then extruded through the slit die in the form of ribbon shape at 3 rpm screw speed under flush mode.

### 4.5. Hot melt extrusion using continuous extruder

For continuous extrusion of biocomposite TPU, spore and TPU powders were premixed at 0 ∼ 1 % (w/w) spore concentration. The mixture powder was introduced into Process 11 (Thermo Fisher Scientific) equipped with a circular die (diameter: 2 mm). All zones except for the feed were set to 135 °C and the mixture was melted, mixed and conveyed at 50 rpm screw speed, which resulted in ∼ 2 min residence time. The composite was extruded into filament shape.

### 4.6. Shear stress calculation

Shear stress during melt extrusion was indirectly calculated from the torque applied to the motors of extruders[9].

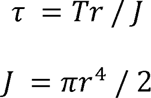

where τ is shear stress, T is applied torque obtained from built in sensors in extruders, r is radius of screws and J is polar moment of inertia.

### 4.7. Injection molding

Biocomposite TPU filament extruded by Process 11 was further processed via injection molding. The filament was introduced into HAAKE^TM^ MiniLab II extruder (Thermo-Fisher Scientific) at 145 °C at 36 rpm under continuous mode. The TPU melt was transferred to an injection cylinder heated at 140 °C. After assembling the cylinder into the HAAKE^TM^ Minilab II ram injection molder (Thermo-Fisher Scientific), a piston purged the polymer melt into the mold at 100 °C under 300 bar pressure for 5 s, followed by post injection under 600 bar pressure for 3 s. The specimens were held in the mold for 1 min and were submerged into a NaCl-saturated ice bath following removal to prevent shrinkage. Biocomposite TPU was injection molded by using a steel mold for fabricating ASTM D638 Type V tensile specimens.

### 4.8. Tensile testing

For the tensile testing of biocomposite TPU prepared using batch extruder, the ribbon-shaped extrudate was tailored into dogbone form by using templated cutting method (overall length: >38□mm; clamping area length: >10□mm; initial distance between grips: ∼18□mm; length of narrow parallel-sided portion: 10□mm; width at ends: 5□mm; width at narrow portion: 2.4□mm; thickness: 0.7□mm). Universal testing machine (Instron 5982, Norwood, MA, USA) equipped with a 100 N load cell was used for tensile testing. Load (N) vs. extension (mm) curves were obtained during stretching of the specimen at 20 mm min^-1^ rate. The data was converted into stress (MPa) vs. strain (-) curves using the following equations.

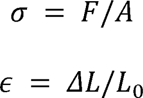

where σ is stress, F is load, A is cross-sectional area of the reduced area of specimen, is strain, ΔL is displacement from the initial specimen length and L_0_ is the initial length of the specimen.

The area under the stress vs. strain curve was calculated based on trapezoid rule to determine the toughness of each specimen. Ultimate tensile stress was obtained from the peak tensile stress during tensile testing. Strain at the moment of fracture was recorded as elongation at break. Young’s modulus was calculated from the slope of the linear part of stress vs. strain curve during the initial stretching of the specimen.

Biocomposite TPUs fabricated using continuous extruder were formulated into dogbone shape by using the aforementioned injection molding method. The specimens were tested using a Shimadzu Universal Testing Machine (AGS-X Series, Shimadzu Scientific Instruments, Columbia, MD, USA) equipped with a 1 kN load cell at an extension rate of 50 mm min^-1^. Same method was followed to calculate tensile properties from raw data.

### 4.9. Spore survivability test

Spores were extracted from the composite by dissolving the polymer component using DMF. In detail, the biocomposite TPU was chopped into small fragments (∼1 x 1 x 1 mm^3^) by using a cutter and soaked into DMF at 40 °C at 150 rpm magnet stirring for 30 min. After centrifugation at 5000 g for 10 min, spore pellets were recovered by discarding the supernatant. Spores were rinsed with fresh DMF one more time, and then suspended in 100 mM PBS pH 7.4.

Spore suspension was serially diluted with 100 mM PBS pH 7.4 prior to colony forming unit (CFU) assay. 100 μL of spore suspension was applied and spread onto LB plates, which were prepared by solidifying 15 mL LB medium supplemented with 15 g L^-1^ agar in each petri dish. After overnight incubation at 37 °C, the number of colonies appearing on the LB plates was counted. It was assumed that each colony was formed from single cells. CFU was obtained by adjusting the number of colonies with the dilution factor and volume of spore suspension applied onto the LB plate. Survivability of spores post processing was calculated by dividing the CFU of spores extracted from biocomposite TPUs by the CFU of spores, which were treated with the same solvent treatment and washing steps of the spore extraction process.

### 4.10. Reprocessing

To evaluate recyclability of TPU and biocomposite TPU, the extrudate was repeatedly processed using a batch extruder. TPU or biocomposite TPU was prepared by following the protocol of batch mode extrusion using HAAKE^TM^ Mini CTW (Thermo Fisher Scientific). The collected extrudates were cut into small pieces below the size of the hopper. 5 g of extrudate was introduced into the extruder and cycled at 135 °C at 36 rpm for 20 min. The recycled TPU or biocomposite TPU was extruded at 3 rpm under flush mode. This process was repeated for up to 4 times of recycling.

### 4.11. Differential scanning calorimetry (DSC)

TA Instruments™ Discovery SDT 650™ (New Castle, DE, USA) was used to determine oxidation induction time (OIT) of TPU and biocomposite TPU. Temperature was ramped to 150 °C under the argon (Ar) environment at 10 °C min^-1^ rate. Ar was replaced with O_2_, followed by isotherm at 150 °C for 2 h.

### 4.12. Storage stability test

TPU and biocomposite TPU were stored in the lab at 23 °C under 60% relative humidity for up to 1 year. Tensile properties and spore survivability were tested after 6 M and 12 M of storage. For tensile testing, templated cutting method was used to prepare dogbone bars. Instron 5982 was used for tensile testing at 20 mm min^-1^ extension rate.

## CRediT authorship contribution statement

**Han Sol Kim:** Writing-original draft, Conceptualization, Investigation, Visualization. **Evan M. White:** Writing-review & editing, Investigation, Project administration. **Grant Crane:** Writing-review & editing, Investigation. **Kush Patel:** Writing-review & editing, Investigation. **Myung Hyun Noh:** Writing-review & editing, Resources. **Md Arifur Rahman:** Writing-review & editing, Resources, Supervision. **Adam M. Feist:** Writing-review & editing, Supervision **Jason J. Locklin:** Writing-review & editing, Project administration, Supervision. **Jonathan K. Pokorski:** Writing-review & editing, Data curation, Supervision, Funding acquisition

## Acknowledgement

This work was supported by BOTTLE^TM^ consortium (# DE-EE0009296) grant funded by the U.S. Department of Energy’s (DOE’s) Office of Energy Efficiency and Renewable Energy (EERE) and Advanced Manufacturing Office (AMO).

