## Supplementary Material for "Scalable Fabrication of a Tough and Recyclable Spore-Bearing Biocomposite Thermoplastic Polyurethane"

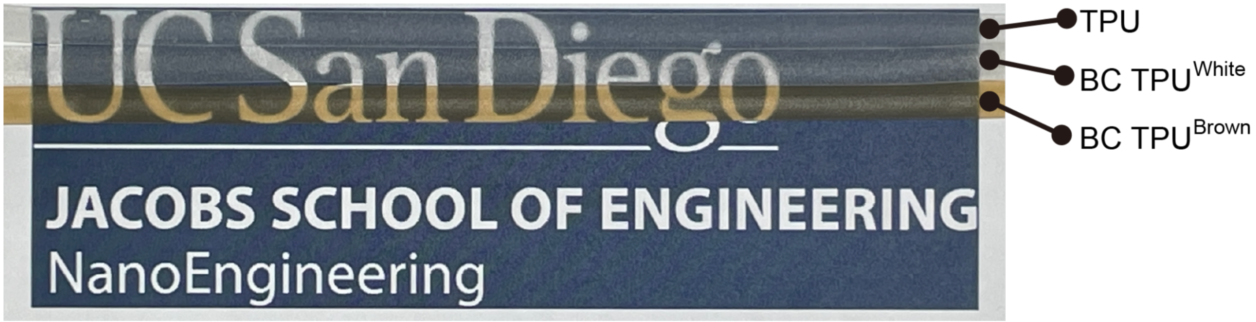


**Fig. S1.** Photograph of TPU and biocomposite (BC) TPUs with 0.5 % (w/w) white or brown spores.

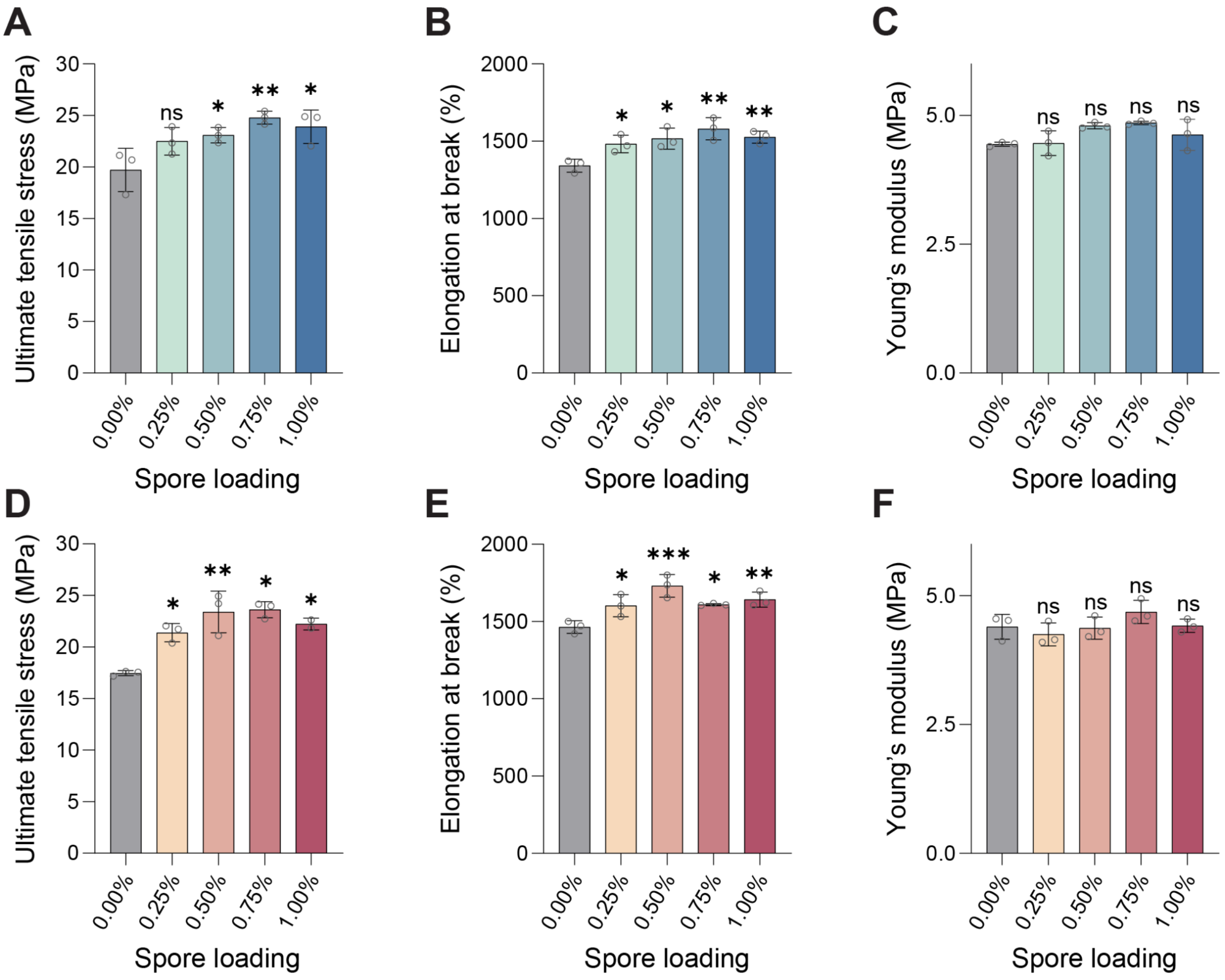


**Fig. S2. Tensile properties of BC TPUs utilizing brown and white spores.** Ultimate tensile stress (A&D), elongation at break (B&E) and Young's modulus (C&F) of BC TPU$^{\text{Brown}}$ (A-C) and BC TPU$^{\text{White}}$ (D-F). One-way ANOVA, followed by a post-hoc test with two-sided Dunnett's multiple comparisons, was used for statistical comparison between TPU (control) and BC TPUs ($n = 3$ per group; ns = not significant, $^{*}P < 0.05$, $^{**}P<0.01$, $^{***}P<0.001$)

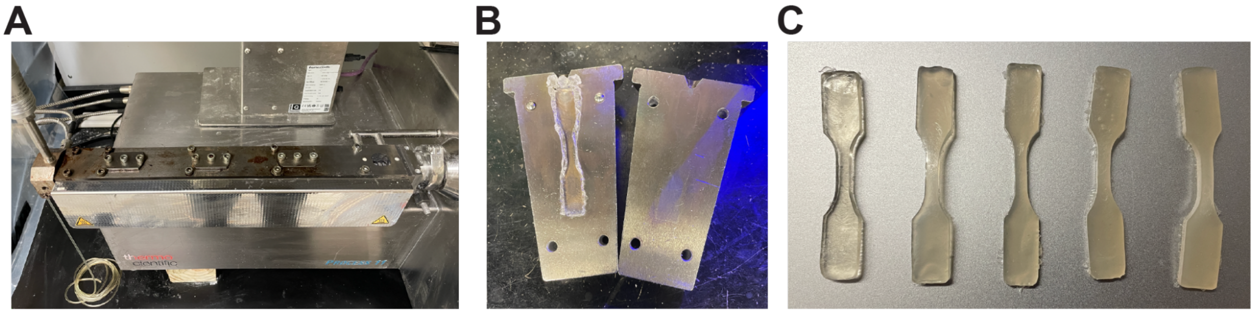


**Fig. S3. Continuous extrusion and subsequent injection molding.** Photographs of continuous extruder (A), ASTM D638 Type V dogbone mold for injection molding (B) and dogbone bars prepared by the combination of extrusion and injection molding for tensile testing (C).

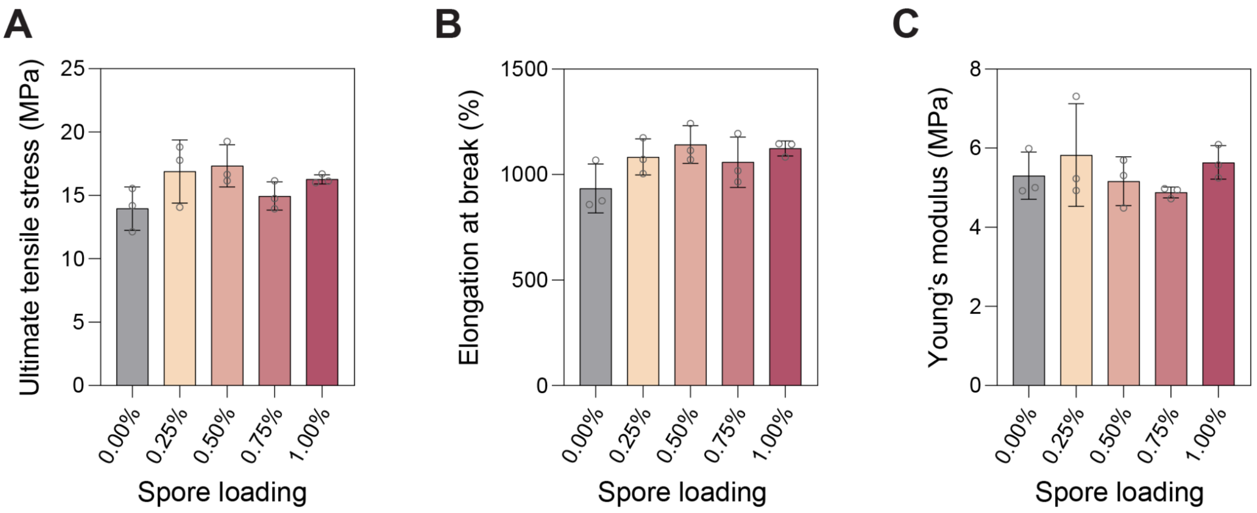


**Fig. S4. Tensile properties of BC TPUs fabricated by Process 11 continuous extruder followed by injection molding.** Ultimate tensile stress (A), elongation at break (B) and Young's modulus of (C) BC TPU$^{White}$ (0.5 % (w/w)).

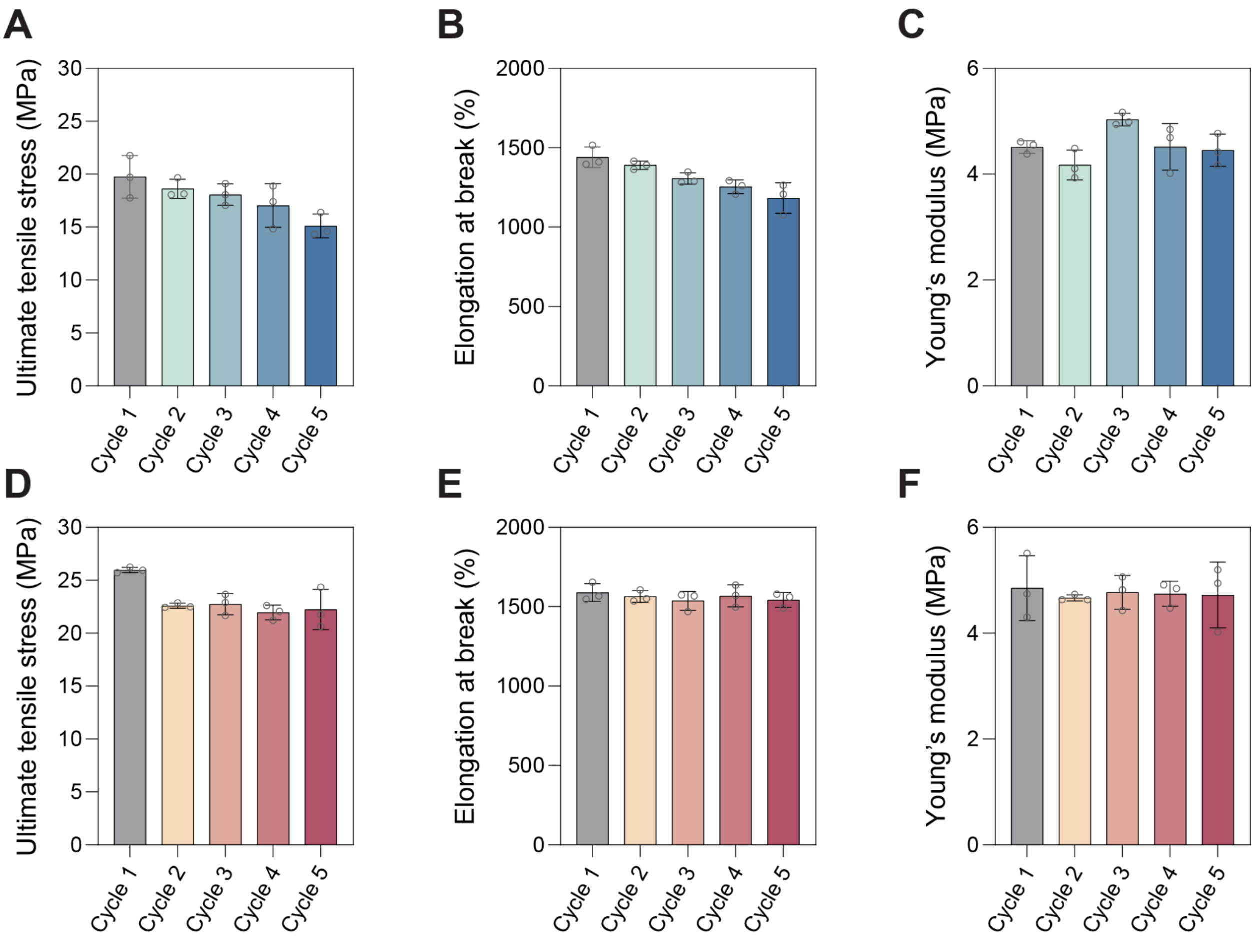


**Fig. S5. Understanding of recyclability of TPU and BC TPUs.** Ultimate tensile stress (A&D), elongation at break (B&E) and Young's modulus (C&F) of reprocessed TPU (A-C) and BC $\text{TPU}^{\text{White}}$ (0.5% (w/w)) (D-F).

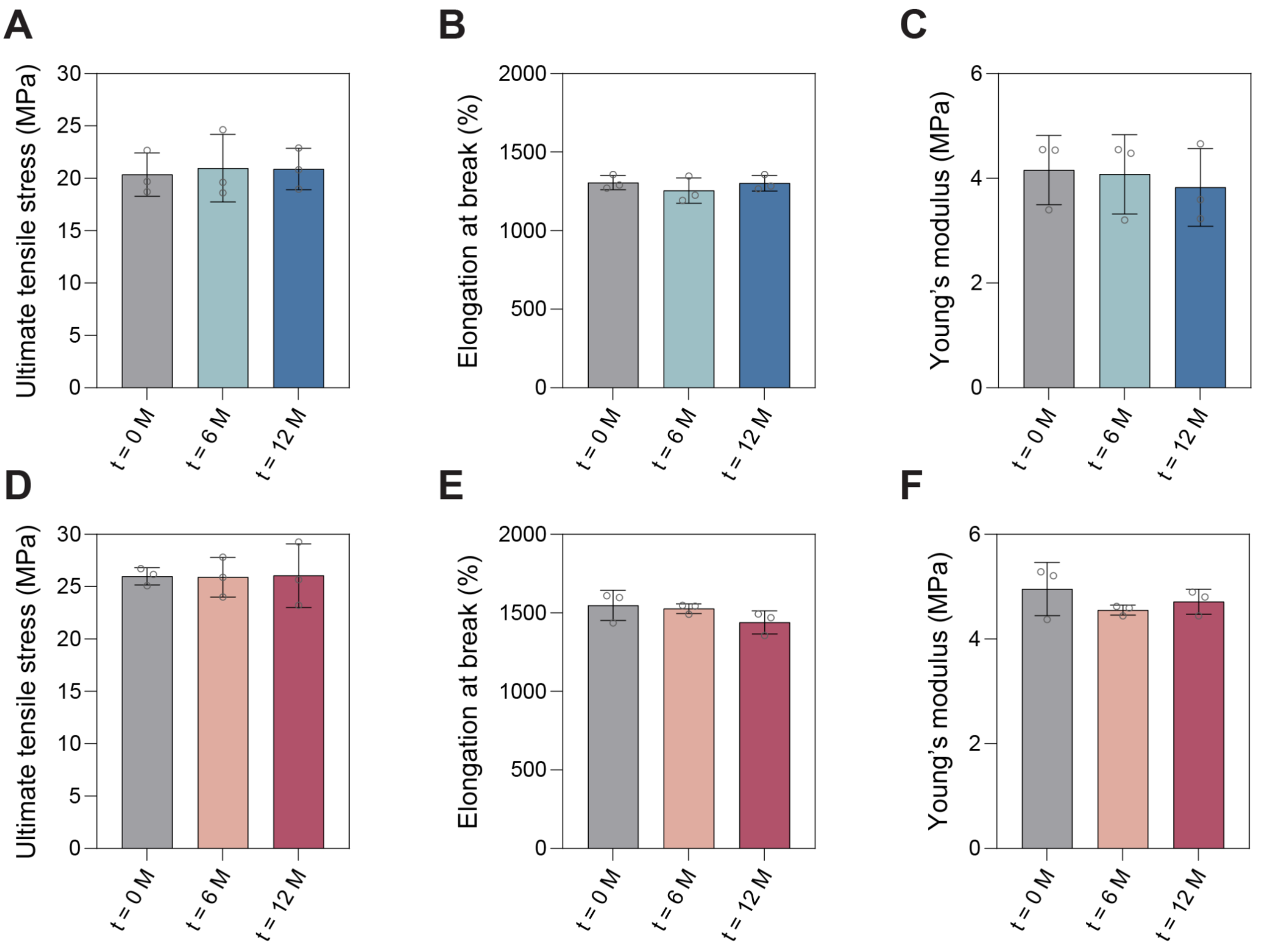


**Fig. S6. Effect of aging (23 °C/60%RH) on tensile properties of TPU and BC TPUs**. Ultimate tensile stress, elongation at break and Young's modulus of TPU (A-C) and BC $TPU^{White}$ with 0.5% (w/w) spore (D-F) during 1 year storage under ambient condition.
